# Automated Image-Based Profiling of Pluripotent Stem Cell Colonies

**DOI:** 10.1101/2025.09.16.676586

**Authors:** Rui Geng, Benjamin L. Kidder

## Abstract

Quantitative image analysis is essential for advancing stem cell biology, developmental studies, and drug discovery, yet most workflows still rely on manual or semi-quantitative scoring that is slow, subjective, and poorly scalable. A major challenge is converting complex colony morphologies into reproducible, high-dimensional datasets. To address this gap, we developed *ColonyQuant*, an open-source platform that integrates automated colony segmentation, alkaline phosphatase (AP) intensity quantification, morphometric profiling, and statistical classification into a single workflow. ColonyQuant computes per-colony functional readouts alongside comprehensive shape descriptors, capturing both staining intensity and structural features in a unified framework. Applied to embryonic stem cells (ESCs) treated with a selective KDM4 histone-demethylase inhibitor, ColonyQuant revealed dose-dependent reductions in colony area and integrated AP signal, together with systematic remodeling of morphometric metrics. Multivariate analyses robustly stratified treatment groups and identified intensity and solidity as principal features capturing dose-dependent colony responses. By transforming subjective scoring into objective, scalable, and biologically interpretable phenotyping, ColonyQuant provides a reproducible platform for stem cell research and high-content screening.

## INTRODUCTION

Embryonic stem cells (ESCs) are central to developmental biology research and hold transformative potential for regenerative medicine due to their unlimited self-renewal capacity and pluripotent ability to generate all somatic cell types^1,2^. The ability to robustly monitor pluripotency status is therefore essential for understanding early cell fate decisions, optimizing differentiation protocols, and screening chemical or genetic perturbations. Among available assays, alkaline phosphatase (AP) staining remains one of the most widely used markers of the undifferentiated ESC state, producing a distinct chromogenic signal that can be rapidly assessed in adherent colony monolayers^3,4^. Despite its ubiquity, AP analysis in most laboratories still depends on manual region-of-interest selection or semi-quantitative visual scoring, approaches that are inherently low-throughput, user-dependent, and prone to bias. These limitations are particularly restrictive in medium-to high-content screening contexts, where large-scale, reproducible, and quantitative assessment of both functional readouts and morphological features is required^5^. Beyond AP intensity, quantitative morphometric descriptors such as area, solidity, and circularity provide complementary measures of colony organization that capture structural phenotypes often overlooked by visual scoring and are linked to cell fate changes and pluripotency.

Automated image-analysis platforms have revolutionized cell profiling by enabling objective, scalable measurements. General-purpose tools such as CellProfiler provide modular pipelines for segmentation and feature extraction^6^, while ImageJ offers an extensible ecosystem of plugins for image processing^7^. Assay-specific scripts such as ColonyArea streamline clonogenic assay readouts^8^, but they typically omit higher-order morphometric descriptors, which are highly informative indicators of self-renewal, early differentiation or subtle perturbation effects and remain underutilized in most workflows. QuPath extends high-throughput analysis to whole-slide imaging and automated immunohistochemistry scoring^9,10^, but its workflows are not optimized for variably shaped stem-cell colonies or integrated morphometric profiling. Interactive machine-learning tools like ilastik enable pixel and object classification without programming^11^, but require downstream scripting to extract AP-specific intensity metrics and complex shape features.

Recent advances in deep-learning–based segmentation, such as Cellpose^12^ and StarDist^13^, deliver robust instance delineation across fluorescence and bright-field modalities, yet they do not directly quantify stain intensities or extract per-colony morphometrics. Classic convolutional architectures such as U-Net^14^ and the DeepCell^15^ framework can achieve high-accuracy segmentation but require extensive annotated training data and remain decoupled from downstream statistical and multivariate analyses. In parallel, high-content morphological profiling studies have shown that rich, multivariate feature sets can capture subtle phenotypes and mechanism-of-action signatures in large-scale screens^16,17^. Despite these advances, there is still no open-source solution that integrates robust colony segmentation with AP-specific intensity quantification, comprehensive morphometric extraction, and downstream statistical modeling into a single, turnkey pipeline optimized for ESC assays.

To close this gap, we developed ColonyQuant, an open-source, modular workflow that unites colony detection, intensity quantification, morphometric profiling, and statistical classification in a single platform. Implemented in Python and R, ColonyQuant is designed for high-throughput reproducibility and seamless extensibility, overcoming the tool fragmentation and parameter sensitivity that limit existing solutions. Colonies are segmented using adaptive Gaussian thresholding and contour detection, enabling accurate delineation across a wide range of staining intensities and colony densities. From these segmentation masks, the pipeline computes per-colony AP intensity metrics and precise area measurements. In parallel, it extracts a comprehensive set of shape descriptors— perimeter, aspect ratio, extent, solidity, equivalent diameter, circularity, and eccentricity— that sensitively capture fine-scale architectural changes. These quantitative features are then integrated into an analysis suite comprising unsupervised (PCA, accelerated t-SNE), supervised (UMAP), discriminant (LDA, PLS-DA), and machine-learning (random forest feature-importance) methods, providing a level of high-resolution phenotyping and classification not available in existing colony-analysis tools. By eliminating manual intervention, ColonyQuant links functional AP readouts directly with rich morphological signatures, enabling unbiased, reproducible discovery of phenotypes.

Complementing its analytical capabilities, ColonyQuant incorporates advanced visualization modules for intuitive, publication-ready data exploration. These include contour-density heatmaps of boundary-pixel frequencies, 2D and 3D density-colored scatterplots of key morphometric features, representative shape-entropy mosaics, and cluster-fraction and inter-group distance heatmaps for quantitative comparisons between conditions. In a demonstration using ESCs treated with vehicle or graded concentrations of a selective KDM4 histone demethylase inhibitor, the pipeline revealed dose-dependent reductions in AP intensity and colony area, accompanied by systematic remodeling of shape metrics. Multivariate classifiers accurately distinguished treatment groups, identifying the most discriminative intensity and morphological features across inhibitor concentrations. The visualization suite further produces violin–box plots, raincloud and ridgeline density plots, z-score heatmaps of group means, radar charts of median feature values, and composite image mosaics—all generated automatically from a single command. Together, these capabilities establish ColonyQuant as the first integrated, assay-optimized, open-source platform for ESC colony analysis.

## RESULTS

### Automated and accurate quantification of AP-stained ESC colonies

To establish a robust baseline for automated colony segmentation, we first applied ColonyQuant to AP-stained ESC colonies. Control ESC colonies displayed compact, well-circumscribed borders in bright-field images, with strong AP chromogenic signal corresponding to undifferentiated states (**Fig. 1A,B**). ColonyQuant segmented colonies with high fidelity, producing binary masks and boundary overlays for each detected object (**Fig. 1C**).

**Figure 1.**
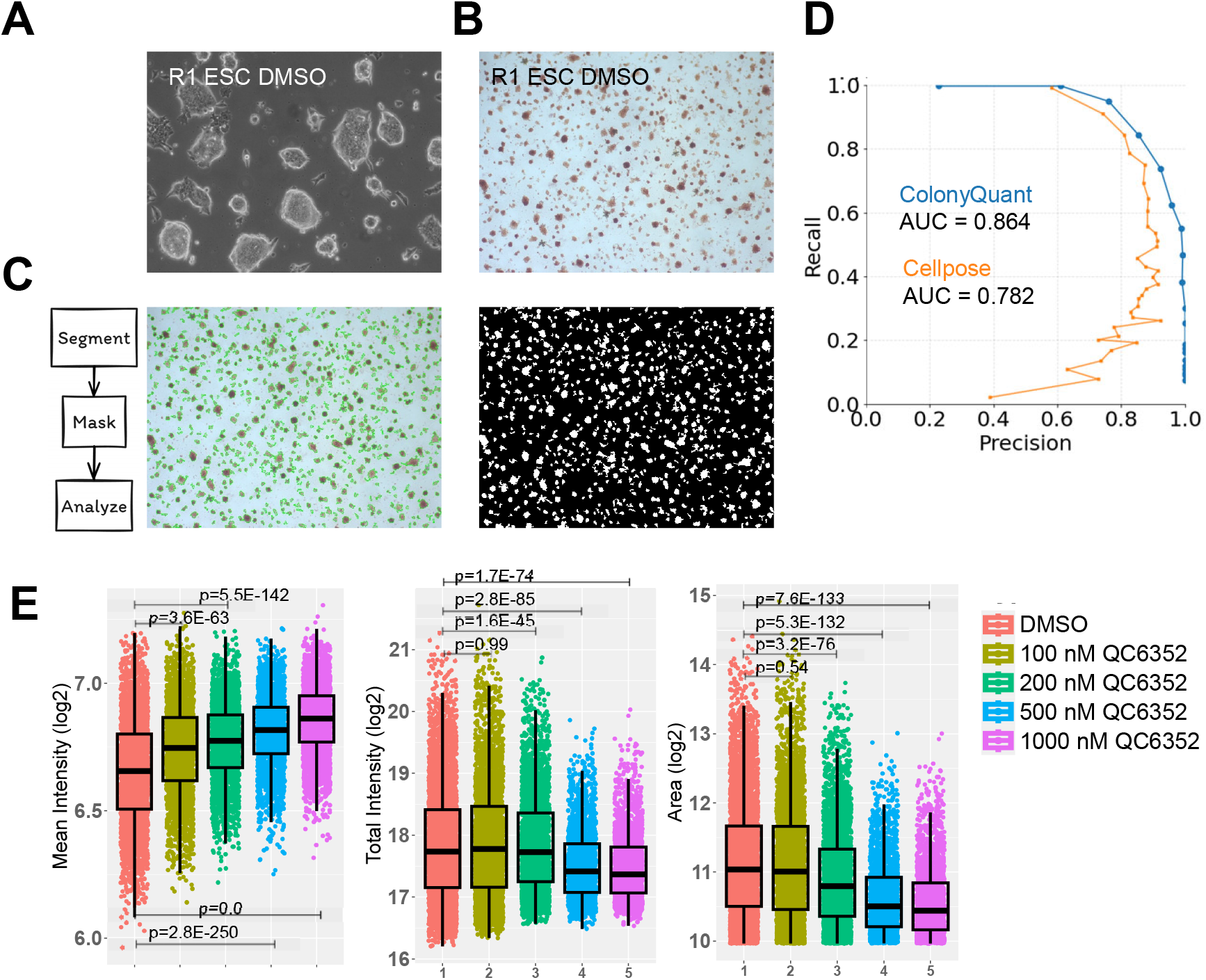
Automated extraction of AP intensity and colony size metrics using ColonyQuant. Example output from ColonyQuant applied to ESC colonies treated with DMSO or increasing concentrations of the KDM4 histone-demethylase inhibitor QC6352. (**A**) Bright-field images of ESC colonies showing compact, well-defined borders. (**B**) AP-stained images highlighting pluripotent colonies with intense chromogenic signal. (**C**) Segmentation masks and overlay boundaries generated by ColonyQuant for each detected colony. (**D**) Precision–recall (PR) analysis against manual annotations comparing ColonyQuant to the best-performing Cellpose variant; ColonyQuant shows a higher PR envelope and larger AUC, indicating better overall detection performance. (**E**) Quantitative metrics derived from segmented colonies: mean AP intensity (left) increased with QC6352 dose, mean colony area (right) decreased, and total AP signal (center), calculated as mean intensity × area, showed the steepest decline between 200 nM and 500 nM, capturing concurrent reductions in colony size and AP output.

Segmentation accuracy was benchmarked against manually annotated ground-truth images and compared directly with the best-performing Cellpose variant. Precision–recall (PR) analysis revealed that ColonyQuant achieved superior performance across the entire curve, with higher recall at matched high precision (**Fig. 1D**). The area under the PR curve (AUPRC) was consistently larger for ColonyQuant, reflecting both improved sensitivity and reduced false-positive rates. Unlike Cellpose, which exhibited marked variability across parameter settings (**Fig. S1**), ColonyQuant’s adaptive Gaussian thresholding and morphological filtering yielded stable, reproducible outputs across images of varying density and staining intensity. These results establish ColonyQuant as a reliable alternative to manual annotation and as a superior solution to widely used deep-learning models for this assay.

### Dose-dependent effects of histone demethylase inhibition on colony growth and AP activity

We next applied ColonyQuant to ESCs treated with graded concentrations of QC6352^18-21^, a selective KDM4 histone-demethylase inhibitor. ColonyQuant quantified dose-dependent reductions in colony size alongside complex changes in AP activity. Mean AP intensity increased progressively from vehicle control to 1000 nM QC6352, whereas mean colony area decreased with dose (**Fig. 1E**). When combined into the integrated metric of total AP signal (mean intensity × area), the steepest decline occurred between 200 nM and 500 nM, capturing the simultaneous suppression of growth. This demonstrates ColonyQuant’s ability to uncover dose–response relationships by integrating multiple morphological and functional features.

### Visualization of spatial and morphological diversity at the population level

To explore population-level heterogeneity, we applied ColonyQuant’s visualization suite to per-colony data. Contour-density maps revealed that control ESC colonies occupied broad peripheral regions of the imaging field, whereas increasing QC6352 concentrations compacted colonies toward the center and reduced overall boundary coverage (**Fig. 2A,B**). Shape-based clustering of Hu-moment descriptors (k = 100) produced representative outline mosaics paired with perimeter-density heatmaps. Controls were dominated by compact, circular colonies, while higher QC6352 doses yielded elongated, fragmented, and irregular outlines, illustrating progressive morphological diversification (**Fig. 2C**).

**Figure 2.**
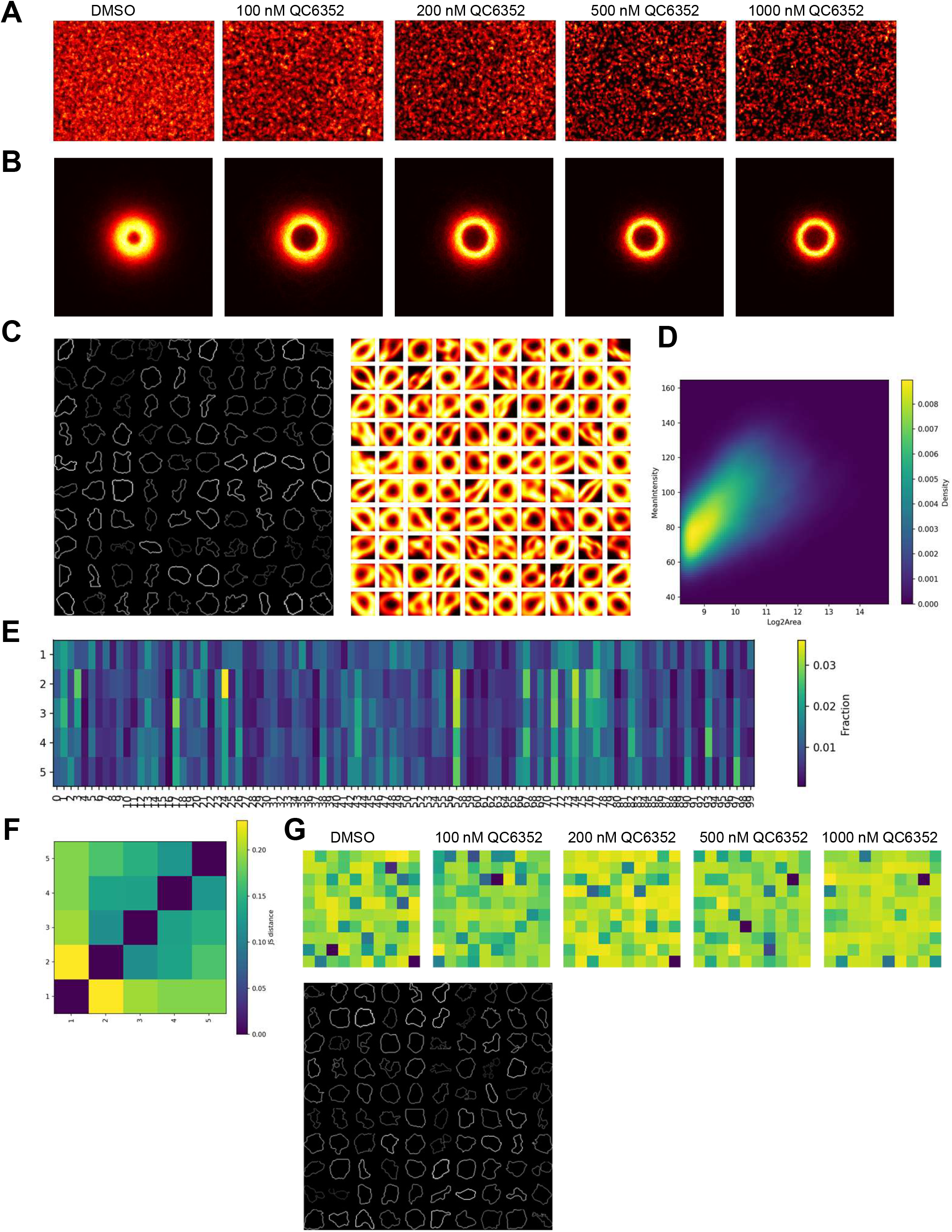
Multi-scale spatial and morphological profiling of ESC colonies. Outputs generated by the ColonyQuant visualization module from AP-stained ESC colonies treated with DMSO or graded concentrations of QC6352. (**A**) Contour-density heatmaps generated by visualize_contours, showing frequency-normalized overlays of all colony boundaries on a fixed 512 × 512 px canvas. (**B**) Individual contour-density heatmaps for each treatment group, with pixel intensity proportional to the frequency of boundary-pixel occupancy. (**C**) Representative-shape mosaics from k-means clustering (k = 100) on Hu-moment descriptors, displaying the contour nearest each cluster centroid (left) and kernel-density maps (σ = 2 px) of all contours within that cluster (right). (**D**) Two-dimensional density heatmap of log_2_(area) versus mean AP intensity (100 × 100 bins; log-transformed densities) summarizing feature-space distributions. (**E**) Jensen–Shannon distance matrix comparing the 100-cluster frequency profiles between treatment groups; darker shading denotes greater morphological divergence. (**F**) Shannon-entropy heatmaps visualizing spatial variability in boundary-pixel occurrence for each treatment group. (**G**) Top row: Shannon-entropy heatmaps of boundary-pixel variability for each condition; bottom: 10×10 mosaic of the 100 highest-entropy contours in the DMSO group, representing the most shape-variable colonies in that condition.

Joint size–intensity distributions further revealed systematic shifts in colony populations. In control samples, colonies clustered along a ridge of large area and moderate AP intensity. Under QC6352, distributions shifted toward smaller colonies (**Fig. 2D**). Cluster-fraction heatmaps demonstrated redistribution of colonies across shape clusters, with Jensen–Shannon distance quantifying minimal divergence between DMSO and 100 nM QC6352, but sharp divergence at 500 and 1000 nM (**Fig. 2E,F**). Boundary entropy analyses showed increasing morphological variability under inhibitor treatment, with high-entropy colonies forming visually distinct subsets absent in controls (**Fig. 2G**). Together, these spatial and structural visualizations highlight ColonyQuant’s capacity to capture subtle and global shifts in colony organization.

### High-resolution morphometric profiling reveals intrinsic variability in ESC architecture

ColonyQuant quantified eight shape descriptors per colony, enabling rigorous characterization of ESC morphologies. Colony area exhibited broad heterogeneity within control populations, spanning nearly an order of magnitude (**Fig. 3A**). Extent values showed that most colonies efficiently filled bounding boxes, although a subset under-filled due to irregular or fragmented edges (**Fig. 3B**). Solidity values confirmed that most colonies were convex, with low-solidity subsets indicating deep indentations or lobulations (**Fig. 3C**).

**Figure 3.**
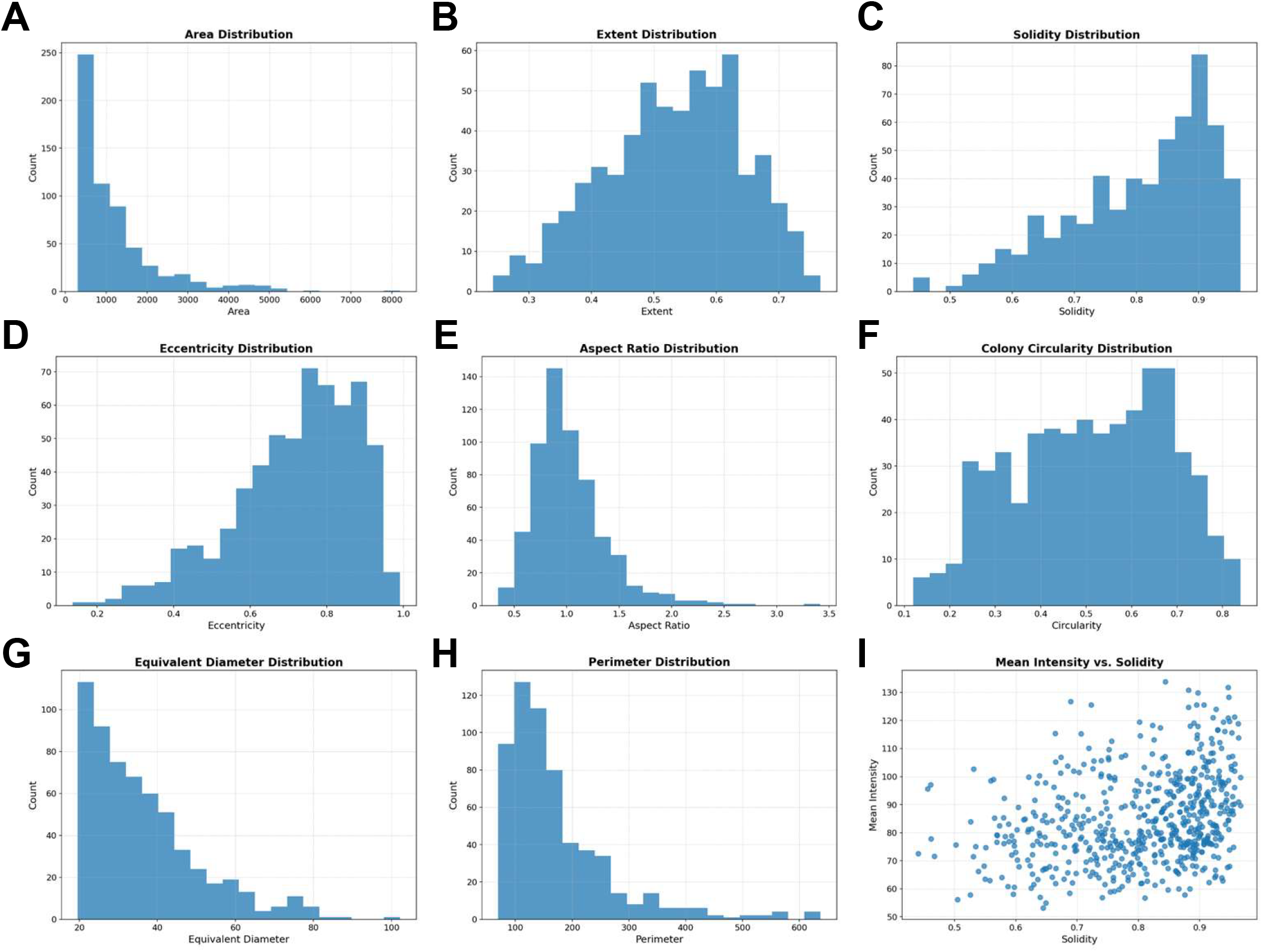
Morphometric diversity and its relationship to AP staining in ESC colonies. Feature distributions and correlations derived from ColonyQuant analysis of AP-stained ESC colonies, highlighting variation in colony architecture and its association with staining intensity. (**A**) Colony area distribution (px^2^) showing broad size variability. (**B**) Extent (mask area / bounding-box area) indicating overall compactness. (**C**) Solidity (mask area / convex-hull area) emphasizing predominantly convex colony shapes. (**D**) Eccentricity (focal-distance / major-axis length) measuring elongation. (**E**) Aspect ratio (major-axis length / minor-axis length) illustrating variation in elongation. (**F**) Circularity (4π·area / perimeter^2^) indicating differences in shape regularity. (**G**) Equivalent diameter (diameter of a circle with the same area as the colony mask) summarizing size in a scale-independent form. (**H**) Perimeter length distribution capturing variation in edge complexity. (**I**) Scatter plot of mean AP staining intensity versus solidity, linking morphological compactness to functional readout.

Other descriptors highlighted further variation: eccentricity and aspect ratio distinguished circular from elongated colonies (**Fig. 3D,E**), circularity quantified boundary smoothness versus complexity (**Fig. 3F**), and equivalent diameter and perimeter captured overall size and edge length (**Fig. 3G,H**). Importantly, mean AP intensity correlated positively with solidity (**Fig. 3I**), suggesting that compact, convex colonies maintain stronger pluripotency marker activity, while irregular outlines may reflect early differentiation or stress responses. These analyses provide quantitative baselines for ESC morphology and reveal functional links between structural compactness and pluripotency state.

### Remodeling of morphological distributions under demethylase inhibition

To illustrate how ColonyQuant profiles entire distributions rather than summary statistics alone, we generated violin–box plots, raincloud plots, ridgeline plots, z-score heatmaps, and radar charts from merged per-colony datasets. Violin plots showed progressive narrowing and downward shifts in mean intensity and colony area under QC6352, while solidity and circularity distributions broadened, reflecting greater morphological heterogeneity (**Fig. 4A**). Raincloud plots emphasized contraction of the long tails for extent and total AP intensity, with higher doses sharply reducing extreme values (**Fig. 4B**). Ridgeline plots highlighted leftward shifts in equivalent diameter and flattening of eccentricity distributions, suggesting altered colony elongation patterns (**Fig. 4C**).

**Figure 4.**
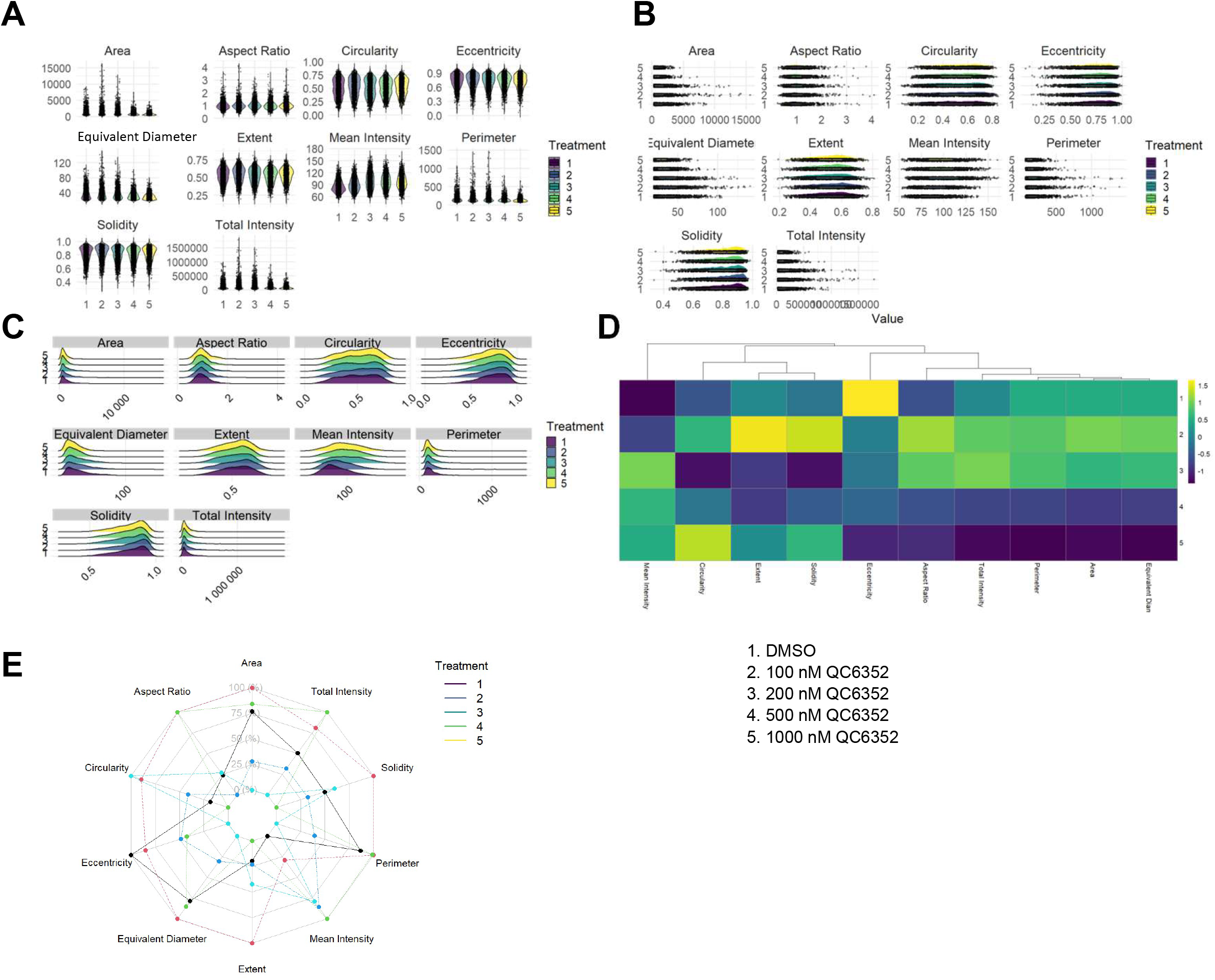
Multi-feature morphological and intensity shifts in ESC colonies under QC6352 treatment. ColonyQuant profiling of ESC colonies exposed to DMSO or 100–1000 nM QC6352 across ten morphometric and intensity features. (**A**) Violin-plus-boxplots showing distributions for Area, Mean Intensity, Extent, Solidity, Circularity, Eccentricity, Aspect Ratio, Equivalent Diameter, Total Intensity, and Perimeter. (**B**) Raincloud plots combining half-eye density curves, boxplots, and individual data points to visualize distribution shape, outliers, and tail behavior. (**C**) Ridgeline density plots (normalized to unit area) stacking features vertically to reveal subtle modal shifts across treatments. (**D**) Hierarchically clustered heatmap of Z-score–normalized group means (rows: treatments; columns: features), with cool tones indicating lower values and warm tones higher values relative to DMSO. Mean Intensity and Solidity show the strongest positive shifts at 1000 nM QC6352, whereas Area and Eccentricity decrease. (**E**) Radar chart of median feature values (scaled to DMSO control) integrating size, intensity, and shape metrics into a phenotypic fingerprint.

A z-score heatmap of group means revealed the most pronounced negative deviations in total AP intensity and solidity at 1000 nM QC6352, whereas aspect ratio and eccentricity shifted positively (**Fig. 4D**). Radar charts condensed these multidimensional changes into distinct phenotypic fingerprints: QC6352 treatment collapsed the size and intensity axes while expanding shape-irregularity spokes (**Fig. 4E**). By automating such multidimensional visualization, ColonyQuant enables rapid generation of phenotypic signatures directly interpretable in terms of underlying biological states.

### Multivariate analysis identifies discriminative morphological and intensity features

Finally, we assessed ColonyQuant’s ability to classify treatment groups using high-dimensional statistical and machine-learning methods. Random forest feature importance ranked mean AP intensity and extent and as the strongest predictors of treatment, with solidity, eccentricity, aspect ratio, and circularity also contributing substantially (**Fig. 5A**). PCA separated DMSO controls from high-dose QC6352 along the first principal component, which explained the majority of variance (**Fig. 5B**). LDA maximized separation, with LD1 alone distinguishing control and treated groups with minimal overlap (**Fig. 5C**). PLS-DA produced clear dose-dependent clustering, with variable importance in projection (VIP) scores again highlighting intensity and solidity as dominant features (**Fig. 5D**).

**Figure 5.**
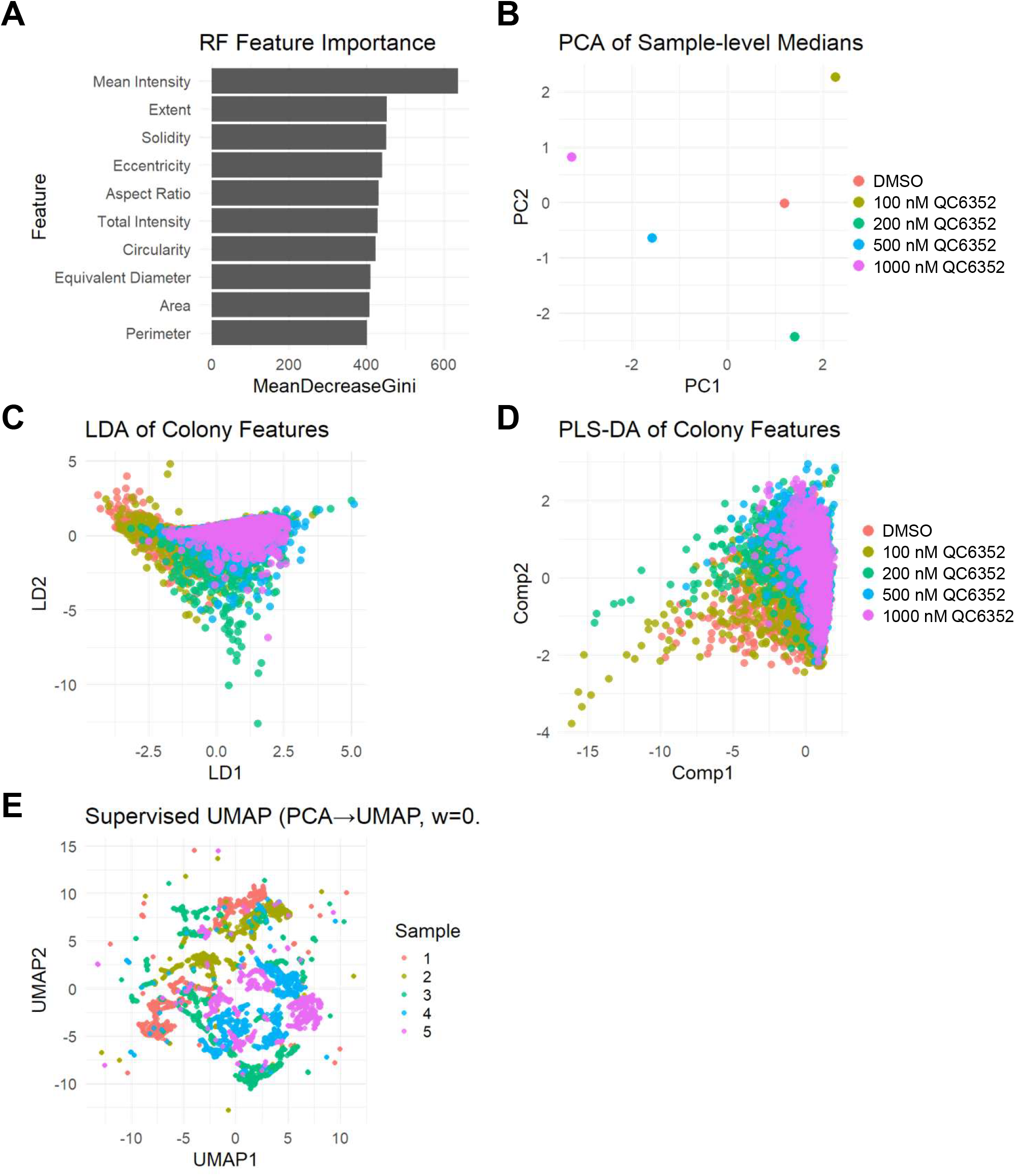
Multivariate classification and embedding reveal QC6352-driven phenotypic separation in ESC colonies. Combined ColonyQuant summary data from DMSO- and QC6352-treated ESC colonies were subjected to a suite of supervised and unsupervised multivariate analyses. (**A**) Random forest feature-importance ranking (MeanDecreaseGini) identifies Mean AP Intensity as the strongest predictor of treatment, followed by Extent, Solidity, and Eccentricity. (**B**) Principal component analysis (PC1 vs. PC2) of sample-level medians separates DMSO controls from high-dose QC6352 primarily along PC1. (**C**) Linear discriminant analysis (LDA) of colony-level features yields near-complete separation of control and treated samples. (**D**) Partial least squares discriminant analysis (PLS-DA; Comp1 vs. Comp2) clusters colonies by QC6352 dose, confirming that both intensity and shape features contribute to separation. (**E**) Supervised UMAP embedding (initialized from PCA, with supervision weight w = 0) projects the top principal components into two dimensions under categorical guidance, producing compact, non-overlapping clusters corresponding to each treatment group.

Supervised UMAP embeddings projected colonies into clusters corresponding to each treatment dose, demonstrating that integrated feature sets capture coherent phenotypic states (**Fig. 5E**). Importantly, these embeddings required no manual intervention, illustrating the accessibility of high-content phenotypic stratification via ColonyQuant. Collectively, these multivariate results underscore how ColonyQuant integrates morphological and functional descriptors to sensitively detect, classify, and interpret treatment-induced phenotypic changes.

## DISCUSSION

### Automated quantification of AP staining and colony morphology enhances objectivity and throughput in stem cell analysis

Alkaline phosphatase (AP) staining remains a cornerstone assay for evaluating pluripotency in ESCs^22^, yet its integration into medium- and high-throughput workflows has been hindered by reliance on manual scoring, subjective interpretation, and inconsistent reproducibility^23^. *ColonyQuant* directly overcomes these limitations by providing a fully automated, end-to-end workflow that unites image preprocessing, colony segmentation, per-object AP intensity quantification, and comprehensive morphometric profiling. By standardizing every processing step, from background correction to high-dimensional visualization, the pipeline minimizes operator bias and ensures reproducibility across datasets, laboratories, and imaging modalities.

Its modular architecture is designed for scalability: individual components can be run independently or chained together, enabling adaptation to different experimental designs, assay formats, and staining types. The same workflow that processes a small, single-condition experiment can be applied—without modification—to hundreds of images spanning multiple treatment groups, producing ready-to-analyze summary tables and publication-quality figures in minutes. By embedding unsupervised and supervised multivariate analysis alongside advanced visualization, *ColonyQuant* enables researchers to move seamlessly from raw images to interpretable phenotypic profiles without custom coding. This combination of automation, reproducibility, and analytical depth makes the tool ideally suited for screening applications in developmental biology, regenerative medicine, and other contexts where objective, quantitative colony assessment is essential.

### Morphometric features as quantitative surrogates of pluripotency state

Beyond intensity measurements, we extract a comprehensive panel of shape descriptors, extent, solidity, circularity, eccentricity, aspect ratio, that provide orthogonal information about colony architecture. Morphological metrics have long been recognized as sensitive indicators of stem cell state^24-28^. In our analysis, solidity and circularity correlate positively with AP intensity, indicating that colonies with compact, convex geometries are more likely to maintain pluripotency. Under QC6352 treatment, both AP intensity and morphological compactness increase, consistent with preservation or reinforcement of cohesive colony architecture. These morphological signatures may reflect structural stabilization that accompanies enhanced pluripotency marker expression, potentially preceding or paralleling transcriptional reprogramming^29-31^. Thus, shape-based descriptors extracted by our pipeline can serve as quantitative, non-invasive proxies for pluripotency^24,25, 27, 32, 33^.

### Multivariate embeddings enable high-dimensional phenotypic discrimination

High-content morphological profiling gains considerable power from multivariate embedding approaches that condense complex feature sets into interpretable low-dimensional spaces while preserving biologically relevant structures^17,34^. Applying PCA, LDA, PLS-DA, and supervised UMAP to integrated AP intensity and morphometric descriptors revealed clear separation between DMSO and QC6352 treatment groups, highlighting the utility of combining functional and structural metrics for phenotype stratification^16,35^. Random forest analysis identified AP intensity, solidity, and colony area as the most discriminative features, in agreement with evidence that chromatin regulators are major determinants of ESC colony architecture and proliferation. Loss of H3K4 demethylases (KDM1A, KDM5B) disrupts compaction and self-renewal, while loss of H3K9 demethylases (JMJD1A, JMJD2C) or gain of H3K9 methyltransferase activity (SETDB1) impairs cohesion and triggers cell-cycle arrest^36-41^. Similarly, perturbation of chromatin-remodeling complexes such as Tip60–p400 abolishes AP staining and distorts colony shape^40^. These parallels reinforce that random forest–derived feature importance can effectively capture epigenetic regulation of colony organization.

### Comparison with existing image analysis tools

Several existing platforms address aspects of AP quantification but fall short of delivering a fully integrated, stain-specific solution. ImageJ offers broad image-processing capabilities but relies on extensive manual parameter tuning and lacks built-in statistical classification^42^. CellProfiler enables flexible, modular pipelines but often demands scripting expertise and can be sensitive to variable staining intensities or low-contrast images^6,43^. QuPath is optimized for histopathology and whole-slide imaging^9^. Moreover, these platforms do not natively provide per-colony morphometric analyses, limiting their ability to capture structural descriptors that are increasingly recognized as critical correlates of cell fate and pluripotency.

Deep learning-based frameworks such as Cellpose^12^ and DeepCell^15^ represent advances in general-purpose segmentation yet they require annotated training datasets and are not inherently designed for stain-specific intensity measurements. In contrast, our pipeline leverages classical computer vision, adaptive thresholding and contour detection, combined with systematic post-processing to achieve high precision and recall with minimal parameter adjustment and no need for labeled training data. This design provides a turnkey, reproducible workflow tailored specifically for AP-stained colony analysis. Benchmarking further demonstrated that ColonyQuant outperformed Cellpose in accurately identifying ESC colonies, reflecting the advantages of a stain-specific, morphology-aware workflow.

### Scalability, generalizability, and accessibility

The pipeline’s open-source architecture and compatibility with standard image formats make it broadly accessible. Its modular design enables straightforward adaptation to other chromogenic assays (e.g., β-galactosidase, X-gal), nuclear stains, or label-free contrast modalities such as phase contrast and differential interference contrast (DIC). Integrated statistical classification and visualization modules allow users to convert raw image data into publication-ready figures rapidly, supporting transparent, reproducible, and quantitative reporting. While our dataset was generated from ESCs, the workflow is readily applicable to other pluripotent stem cell systems—including naïve and primed human ESCs, induced pluripotent stem cells (iPSCs), and organoid-forming progenitors— expanding its utility beyond the model described here.

### Biological insights into histone demethylase inhibition

Our quantitative analysis revealed a dose-dependent increase in AP intensity and decreased colony area following QC6352 treatment, providing new evidence for the compound’s capacity to enhance colony compaction. QC6352 is a potent, selective inhibitor of the KDM4 family (KDM4A–D), which modulates H3K9 and H3K36 methylation in ESCs^18,21^.

Genetic or pharmacologic loss of JMJD2/KDM4 demethylases disrupts ESC self-renewal and proliferation: conditional knockout of Jmjd2a and Jmjd2c causes spontaneous differentiation, colony decompaction, and cell-cycle arrest^37,44^. Jmjd2c is required for stabilizing mediator–cohesin assemblies at lineage-specific enhancers, ensuring proper colony organization and transcriptional programs^45^. In addition, KDM4A and KDM4C drive ESC differentiation into endothelial lineages, linking demethylase activity to overt morphological transitions^46^. The alignment between these known phenotypic consequences and the top features identified by our random forest classifiers, AP intensity, colony solidity, and colony area, demonstrates that high-content imaging combined with machine learning provides a sensitive, non-invasive approach for monitoring epigenetic perturbations in live ESC colonies.

## Conclusion

We present a comprehensive, open-source platform for quantitative AP staining analysis in ESC colonies. By integrating precise image segmentation, multidimensional feature extraction, robust statistical modeling, and high-quality visualization, our workflow delivers a powerful solution for high-content phenotypic screening. Its scalability, adaptability, and ease of use position it as a valuable resource for developmental biology and regenerative medicine, while its modular framework offers a template for extending automated morphological profiling to a wide range of assays and cell systems.

## MATERIALS & METHODS

### ESC culture

R1 mouse embryonic stem cells (ESCs) were cultured under feeder-free conditions on 0.1% gelatin-coated dishes in standard ESC medium composed of Dulbecco’s Modified Eagle Medium (DMEM) supplemented with 15% fetal bovine serum (FBS), 2 mM L-glutamine, 0.1 mM non-essential amino acids, 0.1 mM β-mercaptoethanol, 100 U/mL penicillin, 100 µg/mL streptomycin, and 1,000 U/mL leukemia inhibitory factor (LIF; ESGRO)^39,47^. Cultures were maintained at 37°C in a humidified atmosphere with 5% CO_2_ and passaged every 2–3 days using 0.05% trypsin-EDTA.

To assess the effect of chromatin modulation on pluripotency, cells were treated with DMSO (vehicle) or the KDM4 inhibitor QC6352 (MedChemExpress) at final concentrations of 100, 200, 500, or 1000 nM. At least three independent biological replicates were performed per treatment group. For all conditions, cell seeding density and media volume were standardized to ensure comparability. Cells were passaged routinely by washing with PBS and dissociating with trypsin, using serological pipettes (sc-200279, sc-200281).

### Alkaline phosphatase staining and imaging

Following drug treatment, cells were washed twice with PBS and fixed in 4% paraformaldehyde (PFA) for 1 minute at room temperature. Fixed cells were rinsed with PBS and stained using the Vector Red Alkaline Phosphatase Substrate Kit II (Vector Laboratories) according to the manufacturer’s protocol. Bright-field images were acquired using an inverted microscope. To reduce spatial sampling bias, image fields were randomly selected from each well, with non-overlapping fields captured per replicate.

### Image processing, feature extraction, and statistical analysis

Image analysis was initiated via the `colonyquant quantitate` command-line entry point, which invokes a parser that calls the `process_quantitation()` function. This function, in turn, runs the core `process_image()` routine on each image in the specified directory. The pipeline executes in six stages, each responsible for discrete processing, quality control, or analysis task.

### Preprocessing & initial segmentation

`process_image()` begins by applying rolling-ball background subtraction (radius = 50 px) to bright-field images to correct uneven illumination. Images are then converted to grayscale and processed using adaptive Gaussian thresholding (block size = 51 px, offset = 5) to generate binary masks. To remove noise, two iterations of morphological closing and one iteration of opening are applied using a 5 × 5 elliptical kernel. External contours are extracted, and only regions with an area ≥ 300 px^2^ (user-adjustable) are retained as valid colonies.

### AP intensity quantification

For each retained contour, a filled mask is applied to the red channel (corresponding to alkaline phosphatase staining). Pixel intensities *I* are inverted (255 − *I*) so that values scale directly with staining strength. The mean and total inverted intensities are calculated for each colony, and per-image results are saved alongside binary masks and boundary-overlay visualizations.

### Benchmarking against manual annotations

Manual ImageJ point ROIs placed at colony centroids served as ground-truth positives. A predicted colony (binary mask) was scored a true positive (TP) if the ground-truth centroid fell inside the mask after a small tolerance dilation of the centroid (disk radius 2 px). Unmatched predictions were false positives (FP) and unmatched centroids were false negatives (FN). For each operating point we computed precision =TP/(TP+FP) and recall =TP/(TP+FN). Curves were ordered by recall and the area under the PR curve (AUC) was computed by trapezoidal integration. Metrics were aggregated across all annotated fields.

### ColonyQuant PR curve

ColonyQuant segmentations were generated once per image. The operating point was swept by varying the minimum colony area over a broad range (≈ 0–2600 px^2^) to trace the PR curve. The nominal operating point referenced in the Results (∼ 300 px^2^) achieved > 90 % precision and > 90 % recall on the validation set.

### Cellpose baselines and definition of sweep

We benchmarked Cellpose/Cellpose-SAM across a parameter sweep, i.e., running Cellpose on each image over a small grid designed to be recall-permissive: flow_threshold {0.0, 0.4, 0.8, 1.0, 1.5, 2.0}; cellprob_threshold {−1.0, −0.75, −0.5, −0.25, 0.0, 0.05, 0.1}; tile_norm_blocksize {0, 100, 200, 300}; diameter {0, 60, 90, 120} (0 = auto); and min_size {0, 16, 32}. Each combination defines a Cellpose variant and produced a labeled mask and a cell-probability map per image. For each variant we then performed a threshold sweep by scanning the probability threshold (T ≈ 0.01–0.99) on the probability map to trace its PR curve; connected components above threshold defined predicted colonies. (A post-hoc minimum-area filter was available but is not used in **Fig. S1**.). Among all variant-level probability-sweep PR curves, we selected the single best Cellpose variant by maximum PR-AUC (ties broken by higher recall at ≥ 0.90 precision). **Fig. 1D** plots this best Cellpose curve head-to-head with the ColonyQuant PR curve.

### Morphometric descriptor extraction

Using the `compute_shape_features()` function, eight geometric descriptors are computed for each colony: area, perimeter, aspect ratio, extent, solidity (area / convex hull area), equivalent diameter, circularity (4πA/P^2^), and eccentricity (from ellipse fitting). Mean and total AP intensities from the previous step are appended, and the combined feature set is written to per-image CSV summaries.

### Batch orchestration and aggregation

The pipeline automatically processes all images in the input directory, grouping files by filename prefix, creating output folders, and saving masks, boundary overlays, and per-image summaries. Adjustable parameters, including threshold block size, minimum area, and kernel size, are stored in a single `pipeline_parameters.xlsx` file for rapid tuning.

### Feature-level visualization

Merged `<group>_summary.csv` files are used to compute log_2_(area), identify numeric features, and generate: (i) 3D density-colored scatter plots of any three features; (ii) 2D density heatmaps for any two features; (iii) mosaic grids of representative colony outlines (via k-means clustering on Hu moments); and (iv) heatmap mosaics of contour-density maps. Visualization functions ensure consistent styling, axis labeling, and export of publication-ready figures.

### Quality control and performance assessment

Per-image summaries are aggregated into treatment-level datasets, and bar plots of AP intensity and colony area are generated for rapid assessment of replicate consistency. Segmentation performance is evaluated by comparing automated masks to manually annotated ground-truth masks across a range of minimum-area thresholds. Precision, recall, F_1_-score, and intersection-over-union (IoU) are calculated for each threshold. At the standard cutoff of 300 px^2^, the pipeline achieves >90% precision and recall with a mean IoU of 0.87, indicating robust sensitivity and specificity in colony detection.

### Contour-density heatmaps

All colony masks for each treatment group were processed by the `visualize_contours(size=512)` function. First, each binary mask’s contour pixels (x_i_,y_i_) were mapped onto a 512 × 512 matrix M, where 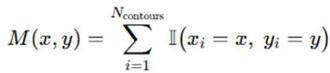. This raw count map was smoothed by convolution with a Gaussian kernel to 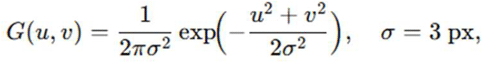 yield *M ′*(*x,y*) = (*M *G*)(*x,y*)

We then linearly normalized M′M’M′ to the unit interval and rendered it with Matplotlib’s “hot” colormap, producing the contour-density heatmaps in Fig. 2A–B.

### 3D Density Scatterplot (Optional Output)

Using the `plot_3d_density()` function, each colony *j* was represented by the feature vector **f**_*j*_ = (log_2_*A*_*j*_,*I*_*j*_,*C*_*j*_), where *A*_*j*_ is area, *I*_*j*_ is mean AP intensity, and *C*_*j*_ is circularity. A Gaussian kernel density estimator in three dimensions computes 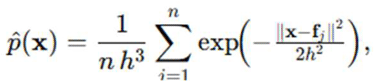 with bandwidth h=0.5. The point cloud is plotted on a 3D Axes (Matplotlib projection=‘3d’), coloring each point by via the viridis colormap. This output appears as colony_3d.png.

### Representative-Shape Mosaics

Contours for each treatment (and separately for the pooled dataset) were encoded by their seven Hu moments [H_1_,…, H_7_]. K-means clustering (k=100) minimized 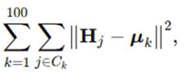 where **μ**_k_ is the centroid of cluster C_k_. For each cluster, the individual contour j∗ with minimal ∥**H**_j∗_−**μ**_k_∥ was rendered as a white outline on a black canvas. All other contours in *C*_*k*_ were overlaid to form a count map *M*_*k*_, smoothed with a Gaussian kernel (σ = 2 px), normalized, and colormapped (“hot”) to generate the density mosaic. These appear as the paired tiles in **Fig. 2C**.

### Two-Dimensional Density Heatmaps

Joint size–intensity distributions were plotted using `plot_2d_heatmap()`. Colony measurements (*x*_*j*_,*y*_*j*_)=(log_2_*A*_*j*_,*Ij*) were binned into a 100×100 grid, with bin counts n_ij_.

Probability densities 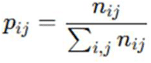 were offset by ε=10^−6^ and log_10_-transformed:

*D*_*ij*_ *=* log10 (*p*_*ij*_ + *ε*) The matrix *D* was smoothed via a 2D Gaussian filter (σ = 1 bin) and displayed, yielding **Fig. 2D**.

### Cluster-Fraction Heatmap and Jensen–Shannon Distance

For each treatment, cluster-fraction vectors p^(g)^=(*p*_1_,…,*p*_100_) were formed, where p_k_ is the fraction of colonies in cluster *k*. Pairwise Jeznsen–Shannon distances were computed as 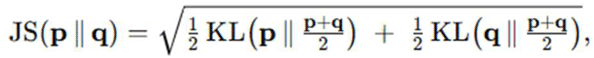 with KL(**p**∥**m**)=∑_k_p_k_log_2_(*p*_*k*_/*m*_*k*_). The resulting 5×5 matrix is rendered as **Fig. 2F**.

### Shannon Entropy Heatmaps and High-Entropy Shape Mosaic

Each treatment’s contour ensemble was rasterized onto a 512×512 canvas and subdivided into a 50×50 grid of bins. For bin (*i,j*), let c_ij_ be the number of contours covering that bin and the total number of colonies; define occupancy probability *p*_*ij*_*=c*_*ij*_*/m*. The Shannon entropy per bin is *H*_*ij*_ = −[*p*_*ij*_log2+(1− *p*_*ij*_) log2 (1− *p*_*ij*_)].

These *H*_ij_ values are normalized to [0–1] and colormapped (viridis), yielding the five heatmaps in the top row of **Fig. 2G**. Separately, each individual contour *j* in the DMSO group was similarly binned to compute a per-colony entropy 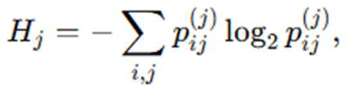 where 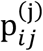 is the fraction of that colony’s boundary pixels in bin (*i,j*). The 100 contours with highest *H*_*j*_ were rendered in a 10×10 mosaic as the bottom panel of **Fig. 2G**.

### Morphological Quantification and Shape Descriptor Analysis

Further morphological characterization was conducted using the morphometrics function, which leverages OpenCV’s contour-analysis and ellipse-fitting routines to quantify a suite of shape descriptors for each segmented colony mask *i*. Colony area *A*_*i*_ was measured as the total number of pixels in the mask, and perimeter *P*_*i*_ as the length of its boundary in pixels.

Extent 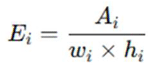 was defined using the mask area and the minimal bounding-box dimensions (width *w*_*i*_, height *h*_*i*_), while solidity 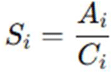 was computed as the ratio of area to the convex-hull area *C*_*i*_.

Ellipse fitting yielded major-axis length ℓ_maj,i_ and minor-axis length ℓ_min,i_, from which we computed: 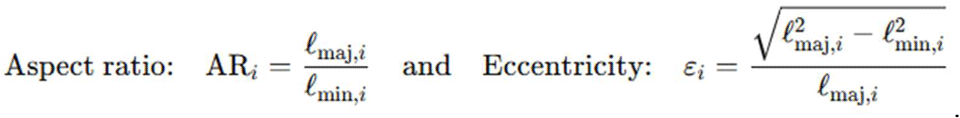.

The equivalent diameter characterizes the diameter of a circle with area *A*_*i*_, 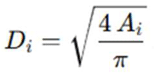 and circularity 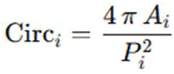 quantifies contour regularity.

Finally, mean AP intensity was obtained by averaging raw green-channel pixel values within each mask: 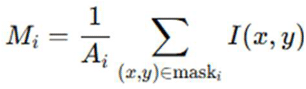 . All metrics were aggregated across colonies, and summary statistics are presented in **Figure 3A–H**. The script also generated histograms for each descriptor and scatter plots, most notably mean AP intensity versus solidity (**Figure 3I**), to reveal relationships between colony shape and pluripotency marker activity.

The summary tables generated were then analyzed using the `feature-distributions` function, which created high-resolution visualizations to illustrate phenotypic changes across treatment conditions. Violin-boxplots revealed full distributions of each feature along with group medians and quartiles. Raincloud plots combined density curves, boxplots, and raw data points to emphasize subtle differences in feature distributions. Ridgeline density plots allowed visualization of modal shifts across treatment groups. To highlight standardized trends, a z-score heatmap of group means was generated. Finally, radar charts synthesized the median feature values across conditions, enabling a multidimensional view of phenotypic shifts induced by QC6352.

### Unsupervised and Supervised Multivariate Analysis and Classification

The final step of the pipeline was conducted with the `comprehensive-analysis` function, which performed unsupervised and supervised multivariate analysis. First, colony-level features were aggregated by computing sample-level medians. Principal component analysis (PCA) and t-distributed stochastic neighbor embedding (t-SNE) were used to explore intrinsic sample structure without labels. Supervised uniform manifold approximation and projection (UMAP) was applied using known treatment conditions to emphasize intergroup separation. Linear discriminant analysis (LDA) and partial least squares discriminant analysis (PLS-DA) were used to identify axes that maximally separated treatment groups. In parallel, a random forest classifier consisting of 500 decision trees was trained to predict treatment category from the full set of intensity and morphometric features. Feature importance scores were ranked using the MeanDecreaseGini index. All statistical analyses were executed with fixed parameters and exported as labeled PNG and PDF plots with high-resolution formatting suitable for publication.

This pipeline produced a complete analysis folder for each dataset, including raw and processed masks, per-colony intensity and shape features, aggregated summary tables, dimensionality reduction plots, classifier output, and full-feature visualizations.

## DECLARATIONS

### ETHICS APPROVAL AND CONSENT TO PARTICIPATE

Not applicable

### CONSENT FOR PUBLICATION

All authors have read and approved the final version of this manuscript.

### Statistics and Reproducibility

All experiments were performed using at least three independent biological replicates per treatment condition (DMSO, 100 nM, 200 nM, 500 nM, and 1000 nM QC6352). For each replicate, non-overlapping image fields were acquired. Image acquisition parameters were held constant, and fields were selected randomly to minimize spatial sampling bias.

Quantitative analysis was conducted using a fully automated, open-source pipeline to ensure reproducibility across replicates. Segmentation accuracy was benchmarked against manually annotated ground truth, yielding an average Intersection-over-Union (IoU) of 0.87. All image processing, feature extraction, and statistical embedding steps were performed with fixed parameters and produced consistent outputs across independent runs. Visualizations and summary statistics were generated using standardized R scripts provided in the analysis module.

## AUTHORS’ CONTRIBUTIONS

B.L.K. conceptualized the study, designed and supported the experimental work, performed data analysis, generated the computational code, and prepared the manuscript. Rui Geng cultured ESCs, performed AP staining, and acquired images.

## ACKNOWLEDGEMENTS

We gratefully acknowledge Zengquan Yang for providing the KDM4 inhibitor QC6352 (MedChemExpress). We also thank the Wayne State University High Performance Computing Grid (https://www.grid.wayne.edu/) for access to the computational resources that made this work possible.

