## Supplemental Figure S1 for "Automated Image-Based Profiling of Pluripotent Stem Cell Colonies"

Running title: Automated ESC Colony Analysis.

*Correspondence:

Benjamin L. Kidder

### Supplemental Figures

**Figure S1. Comparative PR benchmarking of ColonyQuant and Cellpose.**

ColonyQuant precision-recall (PR) curve overlaid with all Cellpose PR curves generated from a parameter sweep. Cellpose was run across a grid of permissive settings—e.g., flow_threshold 0–2.0, cellprob_threshold −1.0 to 0.1, tile_norm_blocksize 0/100/200/300, diameter 0/60/90/120 (0 = auto), min_size 0/16/32. For each resulting variant, a PR curve was obtained by sweeping the probability threshold on its probability map; curves are lightly alpha-blended to show the spread. CQ remains along or above most Cellpose traces at high precision.

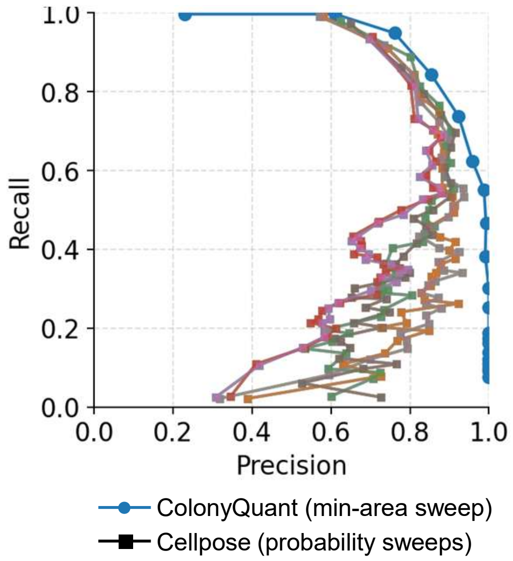
1.0
0.8
0.6
0.4
0.2
0.0
Recall
0.0
0.2
0.4
0.6
0.8
1.0
Precision
ColonyQuant (min-area sweep)
Cellpose (probability sweeps)

**Figure S1**
